# Towards Multi-Brain Decoding in Autism: A Self-Supervised Learning Approach

**DOI:** 10.1101/2025.01.28.635297

**Authors:** Ghazaleh Ranjabaran, Quentin Moreau, Adrien Dubois, Guillaume Dumas

## Abstract

This study introduces a self-supervised learning (SSL) approach to hyperscanning electroencephalography (EEG) data, targeting the identification of autism spectrum condition (ASC) during social interactions. Hyperscanning enables simultaneous recording of neural activity across interacting individuals, offering a novel path for studying brain-to-brain synchrony in ASC. Leveraging a large-scale, single-brain EEG dataset for SSL pretraining, we developed a multi-brain classification model fine-tuned with hyperscanning data from dyadic interactions involving ASC and neurotypical participants. The SSL model demonstrated superior performance (78.13% accuracy) compared to supervised baselines and logistic regression using spectral EEG biomarkers. These results underscore the efficacy of SSL in addressing the challenges of limited labeled data, enhancing EEG-based diagnostic tools for ASC, and advancing research in social neuroscience.

## 1. Introduction

Autism Spectrum Condition (ASC) is a neurodevelopmental disorder marked by difficulties in social communication, repetitive behaviors, and restricted interests (Hodges et al., 2020). Diagnosis typically involves behavioral observations and standardized interviews conducted at various stages of childhood and adolescence (Brihadiswaran et al., 2019; Newschaffer et al., 2007). Diagnosing ASC in adults, however, presents challenges due to a lack of validated tools, as many methods are derived from assessments for children (Lord et al., 2020).

Studying the brain is crucial in ASC research as it provides a direct window into the neural mechanisms underlying social and cognitive differences observed in the condition. Traditional behavioral assessments offer valuable insights but may overlook the biological basis of these challenges, particularly in adults (Murphy et al., 2011). Prior research has revealed alterations in brain structure and function in autistic individuals, such as differences in cortical thickness, connectivity, and activity within social brain networks (Ecker et al., 2015; Uddin et al., 2013). These findings underscore the potential of brain-based approaches to refine diagnosis and deepen understanding of the condition.

A promising diagnostic method, complementary to behavioral assessments, is electroencephalography (EEG), which measures neural activity and offers insight into brain connectivity (Gurau et al., 2017). Resting-state EEG studies in autistic individuals have suggested a U-shaped profile of electrophysiological power alterations (Hull et al., 2018; Milovanovic & Grujicic, 2021), characterized by excessive power in both low-(Hornung et al., 2019) and high-frequency bands (Rojas & Wilson, 2014), abnormal functional connectivity patterns, and enhanced power in the left hemisphere of the brain (Wang et al., 2013). These findings offer valuable neurobiological insights into ASC, particularly with regard to the left-hemisphere asymmetry, which is especially relevant given the language abnormalities commonly associated with the condition (Dawson et al., 1989; Wang et al., 2013).

However, these approaches focus on individuals and task-independent activity, and might not be enough to capture specific features unique to social interactions, such as dynamicity and reciprocity. In the last two decades, researchers have turned to hyperscanning, which monitors brain activity in multiple individuals simultaneously (Montague et al., 2002; Moreau & Dumas, 2021; Novembre & Iannetti, 2021). This approach has unveiled inter-brain correlation (IBC) and inter-brain synchrony (IBS) in a wide range of social scenarios, including imitation (Dumas et al., 2010), mutual gaze (Leong et al., 2017), shared attention (Hirsch et al., 2017), deceptive interactions (Zhang et al., 2017), verbal communication (Hirsch et al., 2018), coordinated activities (Zamm et al., 2018), interpersonal synchronization (Cui et al., 2012), and collaborative tasks (Matusz et al., 2019). These findings challenge the traditional view of the brain as a self-contained system, instead highlighting its role in dynamic, interactive processes (Bottema-Beutel et al., 2019). To advance our understanding and our potential markers of ASC, it is critical to investigate individuals in natural, reciprocal interactions (Dumas, 2022; Nadel & Pezé, 1993). Since social dynamics underpin many psychiatric conditions, hyperscanning offers a promising path to detect and quantify disruptions in social alignment. Difficulties in aligning neural activity with others could serve as a tangible marker of atypical social cognition, offering a more authentic alternative to conventional measures like resting states, but also social perception tasks or Theory of Mind evaluations (Dumas, 2022). Yet, despite its potential, the use of hyperscanning in ASC research is still in its early stages. A pioneering fMRI hyperscanning study by Tanabe et al. (2012) revealed reduced inter-brain synchronization in the right inferior frontal gyrus of ASC-TD (i.e., typically developing) pairs during joint actions, suggesting distinct neural processing in ASC. Similarly, Wang et al. (2020) found that autistic children showed increased frontal cortex synchronization with their parents during cooperative tasks, and this neural synchronization correlated with autism severity. Kruppa et al. (2021) further explored brain-to-brain synchrony in autistic and TD children, finding behavioral but not neural differences, emphasizing the need for deeper investigation.

Hyperscanning with EEG, offering high temporal resolution and real-time insights into social interactions, is particularly well-suited for studying ASC. Nam et al. (2020) reviewed 60 EEG hyperscanning studies, none of which focused on ASC, revealing a significant gap in understanding the social dynamics of this population. This gap is further underscored by the scarcity of ASC-specific research, as noted by Liu and colleagues (2018). Yet recently, it was found that neural synchrony increased during conversations among ASC individuals, with lower synchrony linked to greater social difficulties (Key et al., 2022), as well as differences in inter-brain synchronization between adult ASC-TD pairs during social imitation tasks (Moreau et al., 2023). These findings underscore the potential of EEG hyperscanning to provide valuable insights into the social dynamics and neural underpinnings of ASC individuals.

Despite the great promises of EEG, it is constrained by a low signal-to-noise ratio, non-stationary signals, and substantial inter-subject variability due to individual physiological differences. These challenges make it difficult to extract meaningful patterns from EEG data using traditional statistical methods. To address these issues, computational models are essential for automating pattern recognition, enabling more accurate, reliable, and scalable analysis. Deep learning (DL) models, in particular, are capable of detecting complex patterns in noisy and unstable signals, generalizing across subjects, and automatically extracting relevant features from raw data. This reduces the need for manual feature engineering (Roy et al., 2019). Such capabilities are vital for handling the complexities of EEG data, especially in clinical settings where individual variability is a concern (Wronkiewicz et al., 2015; Zhang et al., 2021). As a result, DL models enhance the ability to detect subtle neural signals, significantly advancing EEG analysis.

DL includes various paradigms, such as supervised, unsupervised, and self-supervised learning. While supervised learning relies on labeled data to train models, self-supervised learning provides an alternative by generating pseudo-labels through data transformations or augmentation (Liu et al., 2021). This allows models to train without the need for external labels, making it particularly advantageous in EEG analysis, where labeling is resource-intensive and requires specialized expertise (Banville et al., 2021; Roy et al., 2019). Given the abundance of unlabeled EEG data, self-supervised learning becomes an invaluable tool, enabling more efficient and scalable analysis. By utilizing self-supervised learning, EEG analysis can become both more efficient and accurate, ultimately enhancing clinical insights. This method reduces the burden of traditional labeling, offering a more scalable solution for advancing data-driven research in the field. Recently, Banville and colleagues (2021) introduced the concept of self-supervised learning for the purpose of acquiring meaningful representations from EEG data. They presented two specific self-supervised learning (SSL) tasks, namely relative positioning (RP) and temporal shuffling (TS) and adapted a third technique called contrastive predictive coding (CPC) (Oord et al., 2019) to be applicable to EEG data.

This study aims to comprehensively analyze hyperscanning EEG data obtained from both autistic and neurotypical participants. The primary objective is to develop a DL model explicitly designed to extract and recognize patterns and relationships within individual EEG signals, employing a self-supervised learning methodology. Building on the work of Bainville and colleagues, we aimed to apply a self-supervised learning approach to the Healthy Brain Network (HBN) and Brain-to-Brain Communication V2 (BBC2) datasets. We first developed a "single-brain model" using SSL on the HBN dataset, which encodes EEG recordings into a concise feature vector. This vector captures characteristics of individual brain activity and extracts distinctive details. For the downstream task, we introduced a "multi-brain model" that simultaneously processes two single-brain models. This model is specifically designed for binary classification, distinguishing interactions involving at least one autistic individual (ASC) from those between two neurotypical individuals (TD). The representations acquired from these EEG signals will play a pivotal role to develop a multi-brain architecture specifically tailored for hyperscanning EEG classification.

## 2. Methods and Material

Self-supervised learning is a dual-stage process, comprising two main components: the pretext task and the downstream task (Jing & Tian, 2019). Pretext tasks serve as the foundational building blocks of SSL, forming a crucial bridge between raw data and the goal of training a model for downstream tasks. In the initial pretext task phase, the model is presented with a set of self-generated challenges or auxiliary objectives. These challenges are carefully designed to encourage the model to extract meaningful and informative features from the unlabeled data. The model learns to uncover patterns, relationships, and representations within the data itself, effectively transforming it into a more structured and informative format.

Once the model has successfully completed the pretext task phase, it has acquired a latent knowledge about the data that is then employed in the downstream task phase, where the model is fine-tuned or transferred to specific target tasks (Liu et al., 2021). These downstream tasks can range from image classification and object detection to natural language understanding and recommendation systems. The model’s pre-trained features, gained through the pretext tasks, enhance its performance on these target tasks.

### 2.1. Datasets

We used two EEG datasets: the Healthy Brain Network (HBN) dataset for the pretext task and the Brain-to-Brain Communication V2 (BBC2) dataset for the downstream task. The HBN dataset (Alexander et al., 2017), from the Child Mind Institute, includes EEG recordings from over 3,000 participants aged 5–21 years, collected using a 128-channel HydroCel Geodesic EEG system at 500 Hz. For this study, we selected resting-state EEG data from 1,000 participants, who alternated between open and closed eyes while fixating on a cross, providing diverse, unlabeled data for self-supervised learning. This large-scale dataset was ideal for training self-supervised models, enabling the extraction of robust feature representations.

For the downstream task, we used the BBC2 dataset (Dumas et al., 2014; Moreau et al., 2023), which includes hyperscanning EEG data from 36 participants in dyads: 9 mixed (TD-ASC) and 9 control (TD-TD). The data, recorded with a 64-channel system at 500 Hz, was collected during three interactive phases: observation of hand gestures, spontaneous imitation, and video-guided imitation. Each phase lasted 1.30 minutes, preceded by a 15-second resting period. This dataset was used to classify dyads based on group composition (TD-TD vs. ASC-TD) and enabled us to fine-tune the pre-trained model from the HBN dataset, assessing its performance in distinguishing neurotypical and autistic interactions.

### 2.2. Preprocessing Pipeline

First, channels that were labeled as ‘bad’, either manually from the BBC2 dataset or by design in the HBN dataset, were interpolated using the spherical spline method to ensure consistency in the data. A notch filter of 50 Hz for the BBC2 and of 60 Hz for the HBN was applied to remove line noise. Using eye EOG channels from BBC2 or proxy EOG channels by creating a bipolar reference from frontal EEG sensors for HBN, we applied Independent Component Analysis (ICA) to remove ocular artifacts, using the extended infomax method, and the ICLabel algorithm to classify components for removal (*MNE-ICALabel — MNE-ICALabel*, n.d.; Pion-Tonachini et al., 2019). Cleaned continuous EEG was then band-pass filtered (0.1 -48Hz) EEG data is typically segmented into epochs, a critical step for accurate IBS computation (Ayrolles et al., 2021). Moreover, preprocessing enhances the performance of machine learning algorithms (Bomatter et al., 2024). Thus, EEG was epoched in constant segments of 1 second, and the remaining noisy epochs were then removed using Autoreject (Jas et al., 2017).

Since the HBN and BBC2 datasets employed different EEG montages (HBN using the 128-channel HydroCel Geodesic Sensor Net (GSN) system (*The Geodesic Sensor Net*, n.d.) and BBC2 following the 64-channel 10-10 Standard System), we aligned the montages for cross-dataset compatibility. The HBN montage was downsampled to match the BBC2 channel count by carefully mapping corresponding channels between the two systems based on established equivalents (<u>EGI Technical Note</u>). This alignment ensured consistency across datasets, enabling effective model training and evaluation. Appendix A provides the full channel mapping details.

### 2.3. Pretext Task

In this study, the model is pre-trained on unlabeled single-brain EEG data using temporal shuffling. We adapted the temporal shuffling (TS) pretask from Banville et al. to the epochs structure by creating labeled samples from the time series *S ∈ R*^*E*^*^✕^*^*C*^ *^✕^*^*T*^, where E is the number of epochs, C the number of EEG channels, and T the number of time samples per epoch. Each sample contains three epochs: two *“anchor”* epochs, *x*_*e*_ and *x*_*e*′′_, and a third epoch, *x*_*e*′_. The index e’ indicates the epoch index at which the window starts in S, and *τ* is the duration of each window.

Since we are dealing with resting state EEG, we establish labels assuming that epochs close in time share similar characteristics. Only windows containing epochs that are less than 3 seconds away are processed.

The *positive context τ*_*pos*_ *∈ N* defines the duration within which two anchor epochs are selected, while the *negative context τ*_*neg*_ *∈ N* provides temporal contrast. Epoch triplets are then classified as either *ordered* (e.g., e < e’ < e’’) or *shuffled* (e.g. e < e’’ < e’), with the label *y*_*i*_ indicating the arrangement. This framework enables the model to learn temporal dependencies within the EEG data, ultimately improving its ability to detect meaningful patterns in hyperscanning tasks.

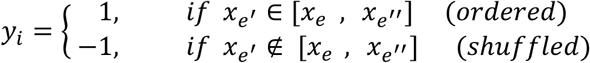

Figure 1A illustrates the sampling process for this pretext task. In each iteration, two anchor epochs, designated in the figure as *"Anchor Epoch A"* and *"Anchor Epoch B"*, are selected, with their temporal distance controlled by a hyperparameter that defines the boundary of the positive context. This positive context is specified as a temporal interval—often represented as a set number of epochs. In the example shown in the figure, the positive context is four epochs (equivalent to four seconds).

**Figure 1.**
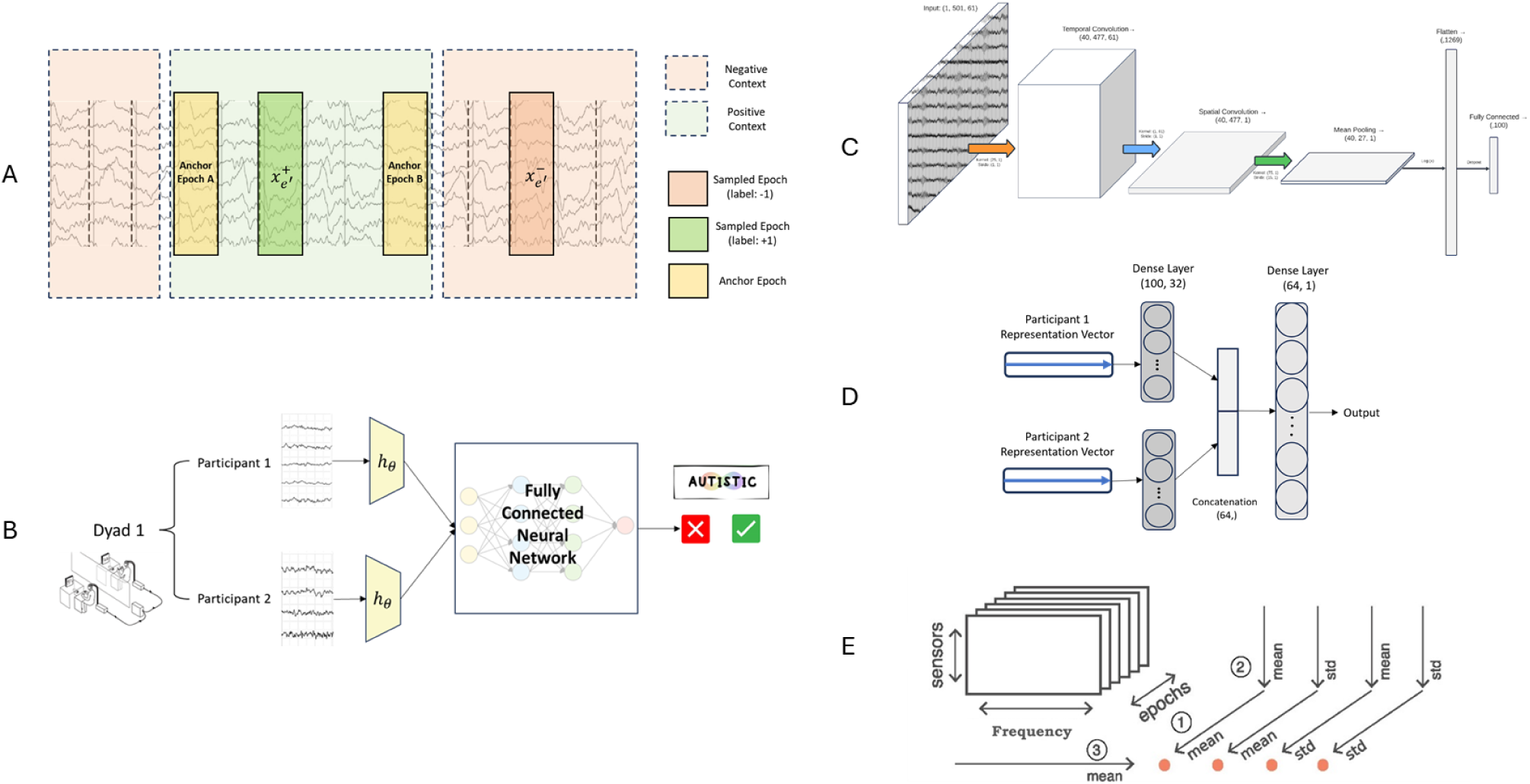
A) Sampling in the TS Pretext Task: Positive context defined by temporal distance between anchor epochs, with positive and negative labels assigned accordingly. B) Downstream Task Model: Visualization of task-specific architecture utilizing pretext embeddings. C) Shallow ConvNet Architecture: Embedder network *h*_*θ*_ for feature extraction. D) Multi-Brain Architecture: Fully connected neural network binary classifier. E) EEG Biomarker Extraction: Methodology based on Denis A. Engemann et al. (2018).

To achieve end-to-end learning in discerning epochs, two essential functions, *h*_*θ*_ and *g*_*TS*_ are required. The function *h*_*θ*_ ∶ *R*^*C*^ ^×*T*^ → *R*^*D*^ serves as a feature extractor with learnable parameters, *θ*, mapping an input epoch from the EEG data into a meaningful *D*-dimensional representation in feature space. The aim is for this feature extractor to capture the underlying structure of the epoched EEG input, enabling effective use of these representations in the downstream task.

Following feature extraction, a contrastive module, *g*_*TS*_, aggregates and evaluates relationships between epochs by contrasting their feature representations. Specifically, for temporal shuffling, the module *g*_*TS*_ ∶ *R*^*D*^ × *R*^*D*^ × *R*^*D*^ → *R*^2*D*^ operates by concatenating the elementwise absolute differences between a test epoch representation, *h*_*θ*_( *x*_*e*′_) and and two anchor epochs, *h*_*θ*_(*x*_*e*_) and *h*_*θ*_(*x*_*e*′′_):

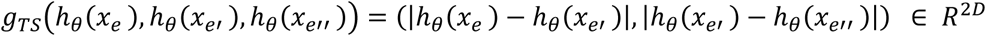

This concatenated output emphasizes the temporal relationships between the epochs, facilitating contrastive learning by focusing on differences between ordered and shuffled samples. A linear context-discriminative model then predicts the target label *y* associated with each epoch triplet by applying binary logistic loss on the outputs of the contrastive module. This setup results in a joint loss function, defined as follows:

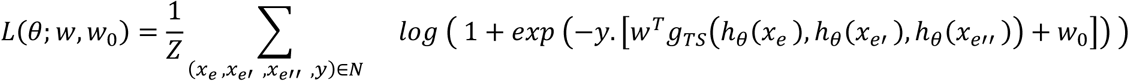

Where Z is a normalization factor, ensuring that the sum is taken over a set N of samples, and (*x*_*e*′_, *x*_*e*_, *x*_*e*′′_, *y*) represents a sample from the set N with *y* indicating the ordered or shuffled label. Parameters *w*^*T*^ and *w*_0_ are those of the linear context-discriminative model, representing the weight vector and bias term, respectively.

By combining the logistic loss with contrastive representations derived from *g*_*TS*_ module, this joint loss function penalizes deviations from the true labels *y*, thereby encouraging the model to learn and generalize temporal dependencies across epochs effectively. Moreover, summing over a set of samples, N, further reinforces the model’s ability to discern temporal patterns across varied EEG epochs.

### 2.4. Downstream Task

Following this pre-training, the model now serves as a feature extractor for a downstream binary classification task, utilizing hyperscanning-EEG data that captures interactions either between neurotypical participants or between neurotypical and autistic individuals.

To adapt the model for this task, we fine-tuned it using a fully connected neural network to classify hyperscanning-EEG features into two categories: autistic or non-autistic individuals (as illustrated in Figure 1B). The model, which has already learned valuable representations in the self-supervised phase, is thus primed for improved performance in the classification task. This end-to-end model, combining the pre-trained self-supervised component with the downstream classification network, ultimately enables the prediction of whether a given interaction, as reflected in the hyperscanning-EEG data, stems from a control dyad or an autistic dyad. This approach underscores the potential of self-supervised learning to enhance model performance in specialized tasks, particularly in classifying neurotypical and autistic interactions.

### 2.5. Model Architecture

The EEG self-supervised learning framework employs embedders as feature extractors to convert each 1-second epoch of data into a 100-element vector that captures the underlying neural activity. These embeddings are subsequently input into a contrastive module to evaluate temporal dependencies between epochs. For feature extraction, a customized Shallow ConvNet (Schirrmeister et al., 2017), originally designed for brain-computer interface applications, is utilized. This architecture comprises a single convolutional layer followed by squaring non-linearity, average pooling, logarithmic non-linearity, and a linear output layer, with batch normalization applied after the convolution. Despite its simplicity, Shallow ConvNet has demonstrated strong performance in EEG-based tasks, including pathology detection (Gemein et al., 2020), and has been effectively used in the temporal shuffling pretext task, as reported by Banville et al. In this study, the model is tailored to process 1-second resting-state EEG epochs with 61 channels sampled at 500 Hz, resulting in an input tensor size of (501 x 61). Adjustments to kernel sizes and dense layer dimensions are made to ensure compatibility with the altered input dimensions, optimizing performance for the task at hand, as illustrated in Figure 1C.

For the downstream task described in Section 2.3, we utilized two pre-trained embedders to generate vectorized representations of hyperscanning EEG data. These representation vectors were subsequently input into a binary classifier, which was implemented as a fully connected neural network. The architecture of the classifier is depicted in Figure 1D.

### 2.6. Baseline Comparison

To evaluate the performance of our model on the downstream task, we compared it against two baseline approaches: (1) a purely supervised model, and (2) a logistic regression classifier using extracted spectral EEG biomarkers. Each baseline approach was designed to highlight different aspects of model performance and compare the effectiveness of the SSL framework with traditional methods.

#### 2.6.1. Supervised Model

In this baseline approach, no pretext training is conducted, meaning the model starts from scratch without any learned representations from the SSL phase. The model is directly trained on the downstream task using the BBC2 dataset, with no prior knowledge or weight adjustments. This comparison evaluates whether the SSL pre-training adds value, as it allows us to assess if the model benefits from prior learning or if training from scratch performs similarly. The goal is to determine the importance of transferring knowledge from the pretext task to the downstream task.

#### 2.6.2. Logistic Regression Classifier with Spectral EEG Biomarkers

In this baseline, we employ a traditional machine learning approach using extracted spectral EEG biomarkers. Following the methodology outlined by Engemann and colleagues (Engemann et al., 2018), we derive 10 spectral features from each participant within a dyad. Four key features are computed for each EEG marker: ’mean-mean’ (mean of the means across epochs and sensors), ’mean-std’ (mean of the standard deviations across epochs and sensors), ’std-mean’ (standard deviation of the means across epochs and sensors), and ’std-std’ (standard deviation of the standard deviations across epochs and sensors), as shown in Figure 1E. These features are calculated for each of the 10 spectral biomarkers and concatenated to form representative feature vectors for each dyad. The resulting feature vectors are then input into a binary logistic regression classifier, which predicts whether the dyad belongs to the TD or ASC group. This traditional machine learning approach is a well-established method for EEG-based classification and serves as a benchmark for comparing the performance of the DL model.

## 3. Results

### 3.1. Pretext Task Model Performance

The pretext model, a single-brain neural network, demonstrated strong feature extraction capabilities on a proxy dataset of 150,123 epoch triplets derived from 1,000 HBN subjects. Each triplet, consisting of two anchor epochs and one sampled epoch, was created using the TS algorithm with a positive context of 10 seconds. Epochs contained 501 time samples across 61 EEG channels. The data was split into training (90%) and evaluation subsets (10%), with the latter further divided into validation (90%) and test (10%) sets. The model was trained for 200 iterations using the Adam optimizer (learning rate: 1^e-5^), a batch size of 128, a dropout rate of 0.4, and 207,021 trainable parameters. While training accuracy reached 100% with near-zero loss, validation accuracy plateaued, indicating overfitting (Figure 2A). Early stopping at iteration 34 mitigated this issue, and test set evaluation, performed with a batch size of 64, yielded an average accuracy of 90% demonstrating the model’s ability to learn and generalize meaningful temporal features from EEG data, as depicted in Figure 2B.

**Figure 2.**
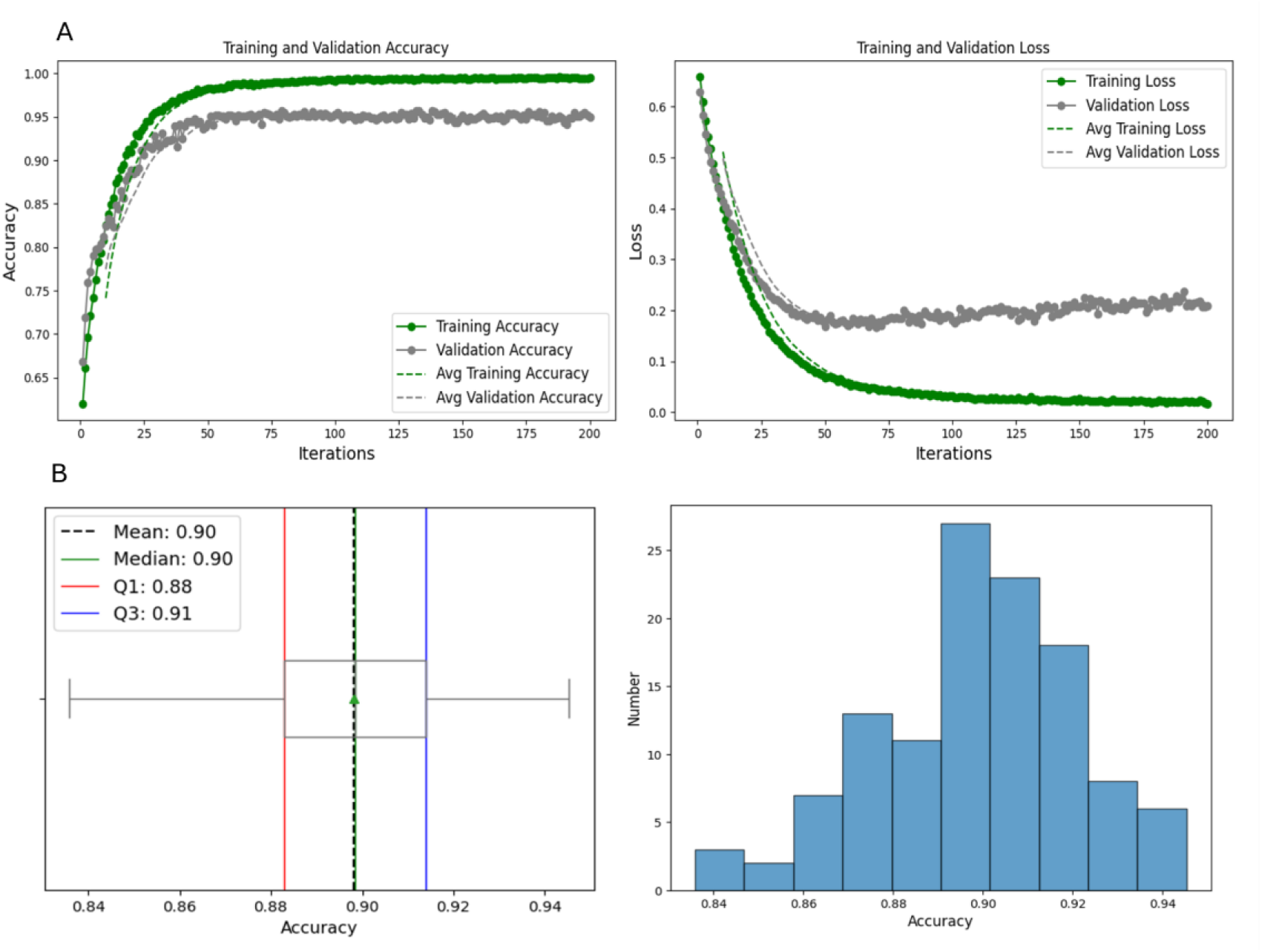
A) Performance of the pretext model with Shallow ConvNet embedder in the training phase. B) Statistical characteristics and accuracy value distribution of the pretext model in the testing phase.

### 3.2. Downstream Task: Performance of the SSL Multi-Brain Model

A downstream binary classification task was performed using hyperscanning EEG data from the BBC2 dataset. The objective was to classify dyadic interactions as either neurotypical (TD) or involving at least one autistic individual (ASC). The dataset comprised 142,120 dyadic epochs, each containing data from 61 EEG channels and 501 time samples, representing a one-second temporal window.

To process these epochs, a multi-brain model was employed, integrating two pre-trained single-brain embedders based on the Shallow ConvNet architecture. The data were divided into training (90%) and evaluation (10%) subsets, consistent with the partitioning strategy used during the pretext task. Model training was conducted using the Adam optimizer with a logarithmic logistic loss function. Hyperparameter tuning was carried out using Optuna (Akiba et al., 2019), a Python-based framework that leverages the Tree-structured Parzen Estimator (TPE). All computations were executed on the Compute Canada Narval Cluster. Among the optimized hyperparameters, dropout emerged as the most influential factor (Figure 3A). The finalized configuration included 79 optimization iterations, a batch size of 8, a learning rate of 9.3e-4, and a dropout rate of 0.38.

**Figure 3.**
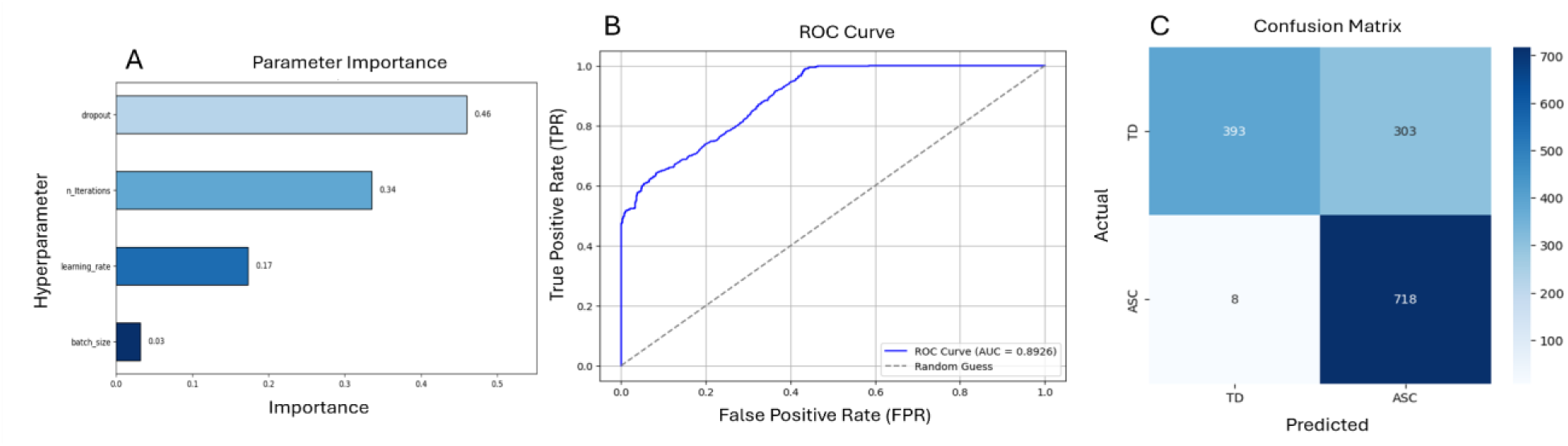
A) Hyperparameter importance in multi-brain model. B) ROC curve, and C) Confusion matrix of SSL Multi-Brain Model.

The multi-brain model achieved a mean test accuracy of 78.13% when evaluated on batches of five unseen dyadic epochs, closely approaching the benchmark accuracy of 79.4% reported by Banville et al. (2021). To further assess performance, additional metrics were calculated, including the ROC curve, binarized predictions at a threshold of 0.5, and a confusion matrix (Figure 3B and 3C).

Key performance metrics underscored the effectiveness of the SSL multi-brain model in distinguishing between neurotypical and ASC interactions. The model achieved a precision of 70.32%, indicating its capacity to accurately identify ASC interactions. Recall (sensitivity) was notably high at 98.90%, reflecting robust detection of ASC cases. The F1 score, a harmonic mean of precision and recall, was 82.20%, highlighting the model’s overall reliability. However, specificity was lower at 56.47%, indicating reduced accuracy in classifying neurotypical interactions. Despite this limitation, the model demonstrated strong overall accuracy (78.13%) and a robust ROC AUC of 0.8926, confirming its strong discriminative ability across both classes.

### 3.3. Baseline Comparisons

To evaluate the impact of SSL pre-training, we conducted baseline experiments using two approaches: a supervised multi-brain model with untrained embedders and a logistic regression classifier trained directly using the BBC2 dataset. The first baseline utilized the same multi-brain architecture as the SSL model but without pre-training the embedders. This supervised model was trained on labeled hyperscanning EEG data from the BBC2 dataset, using identical architecture, dataset splits, hyperparameters, and optimization procedures as the SSL model. Testing on batches of five dyadic epochs resulted in an average accuracy of 52%, slightly above random chance (50%).

Moreover, the model’s performance was further assessed using following metrics: It achieved an ROC AUC of 0.5224, precision of 0.5290, recall of 0.5275, an F1 score of 0.5283, specificity of 0.5101, and overall accuracy of 0.5190. The confusion matrix (Figure 3B) and the ROC curve (Figure 3A) confirmed the model’s limited discriminative capacity, as its results were only marginally better than random chance.

The second baseline involved a logistic regression classifier trained on spectral EEG biomarkers extracted following the methodology outlined in Section 2.4.2. These biomarkers included five primary frequency bands—delta (δ), theta (θ), alpha (α), beta (β), and gamma (γ)—and their normalized versions, resulting in a total of 10 spectral features per participant. Each feature was summarized using four statistical measures—mean-mean, mean-std, std-mean, and std-std— resulting in 40-dimensional feature vectors per participant. Dyadic feature vectors were formed by concatenating the features from both participants, producing 80-dimensional inputs for binary classification.

Despite hyperparameter tuning using <u>GridSearchCV</u>, the logistic regression model achieved a cross-validated accuracy of 50%, a test accuracy of 50%, and an ROC AUC score of 0.50 (Figure 3.A). The classification report revealed that the model failed to predict one of the classes (ASC), with predictions heavily biased toward the majority class (TD). For the class TD, the model achieved a Precision of 0.50, Recall of 1.00, and an F1-Score of 0.67, suggesting that while it successfully identified all instances of TD, half of its TD predictions were incorrect. However, for the ASC class, the model showed poor performance with a Precision and Recall of 0.00, resulting in an F1-Score of 0.00. Confusion matrix confirmed this bias, demonstrating that the handcrafted features were insufficient to enable reliable classification of the dyads (Figure 4D).

**Figure 4.**
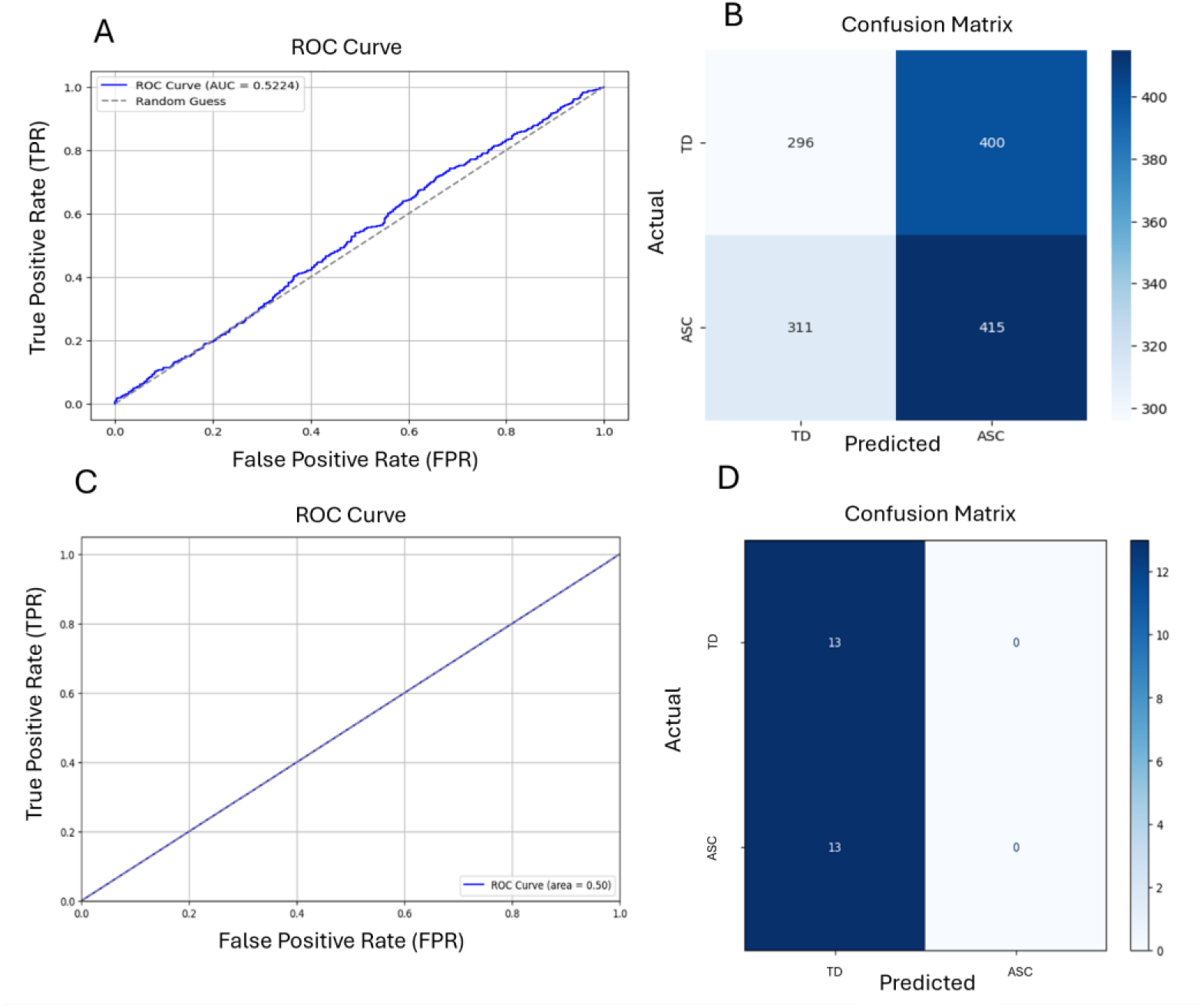
A) ROC curve and B) confusion matrix of the supervised multi-brain model. C) ROC curve and D) confusion matrix of the logistic regression classifier .

### 3.4. SSL Multi-Brain Model Outperforms The Baseline Models

Our analysis reveals significant performance differences among the models (Figure 5). The SSL Multi-Brain model with Shallow ConvNet embedders achieved the highest classification accuracy (78.13%), demonstrating the effectiveness of self-supervised learning in extracting robust features from EEG data. Additionally, it exhibited greater consistency, with a narrow interquartile range and the third quartile aligning closely with the maximum, reinforcing its status as the top performer.

**Figure 5.**
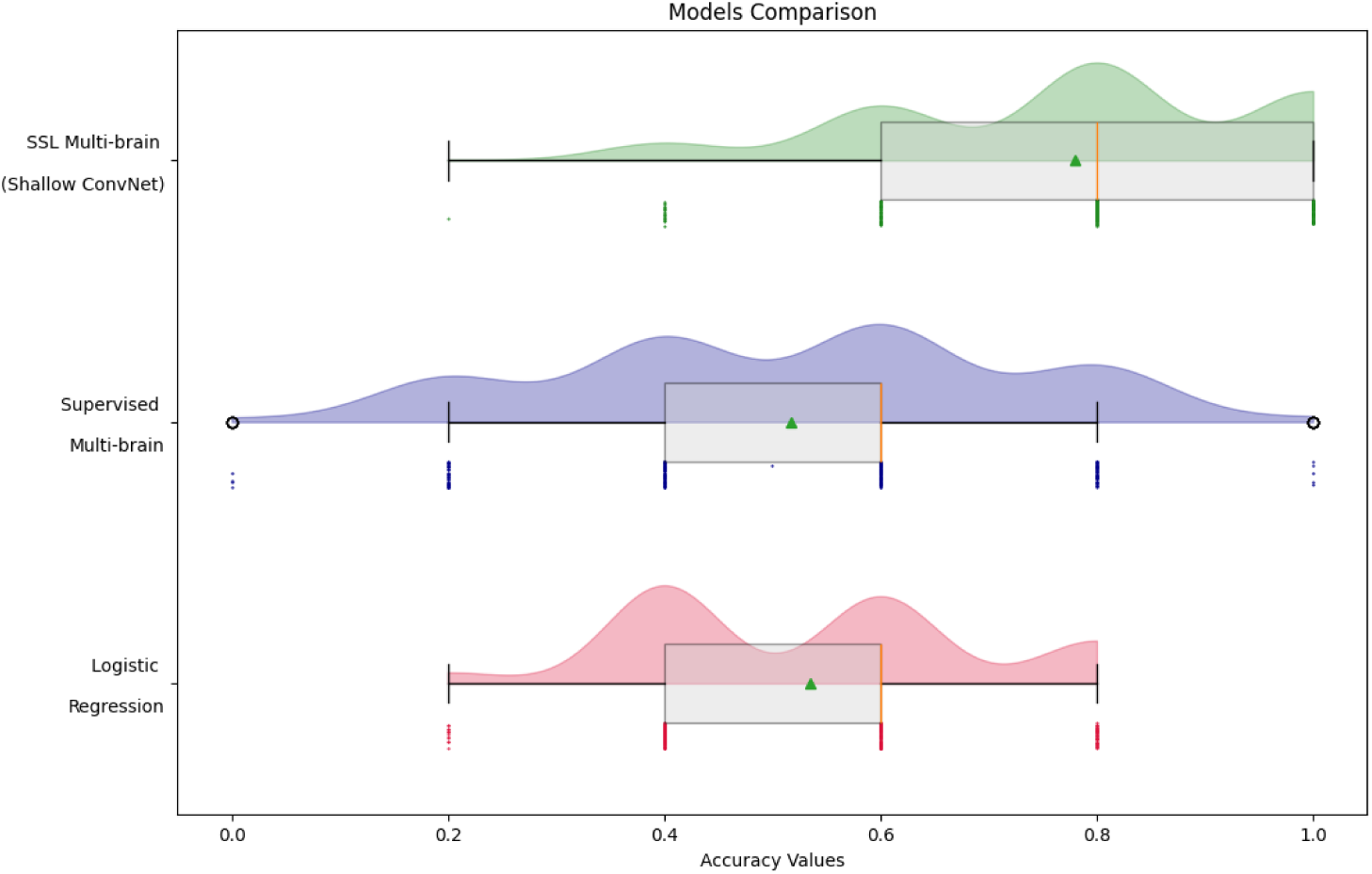
-Comparison of the models’ performances for hyperscanning-EEG classification task.

In contrast, the supervised multi-brain model without pre-training achieved only 52% accuracy, which is comparable to random chance. This highlights the difficulty of training on hyperscanning data without SSL pre-training, which is crucial for learning rich, meaningful feature representations.

For all deep learning models, testing was conducted on 285 dyadic epochs, and accuracy was calculated in batches of five epochs. In contrast, the logistic regression model, which used features extracted from each dyad rather than individual epochs, was evaluated on a testing set of 26 dyads. To ensure comparability of the results, a resampling method was applied to calculate accuracy for these 26 samples in batches of five. The logistic regression model achieved a mean accuracy of 50%, which is equivalent to chance performance, highlighting the challenges of classifying hyperscanning data using traditional machine learning techniques.

In terms of class balance, the SSL model performed well despite the natural class imbalance in the dataset, as it maintained a high recall (98.90%) for detecting ASC interactions. This is critical, as class imbalance often skews model performance, leading to models that favor the majority class (neurotypical). The precision-recall trade-off here highlights the model’s effectiveness in detecting ASC interactions, with a precision of 70.32%. This is a strong performance in identifying ASC cases, even though specificity was lower (56.47%), reflecting the challenge of correctly identifying neurotypical interactions.

The baseline models, on the other hand, struggled with these challenges. The supervised multi-brain model’s precision, recall, and F1 score were low, indicating poor discrimination between the classes. The logistic regression model also demonstrated a heavy bias toward the majority class (neurotypical), with a recall of 1.00 for TD and a recall of 0.00 for ASC, further highlighting its failure to capture the complexity of the data.

These findings demonstrate the superiority of the SSL Multi-Brain model, which not only outperforms traditional and supervised deep learning models but also handles the class imbalance effectively. By leveraging unlabeled data through pretext tasks, SSL extracts more reliable, transferable features, setting a new benchmark for hyperscanning EEG analysis.

## 4. Discussion

In this study, we demonstrated that Self-Supervised Learning (SSL) can be effectively applied to epoched EEG data, yielding strong results in hyperscanning analyses. Specifically, SSL enhanced the performance of binary classification tasks, distinguishing between Autism Spectrum Condition (ASC) and neurotypical (TD) interactions. By leveraging large-scale, unlabelled EEG data, SSL enabled the extraction of meaningful temporal features, which significantly improved the performance of downstream classification tasks. Notably, SSL-based pretraining achieved a mean accuracy of 78.13%, outperforming baseline models, and underscoring SSL’s potential to address the challenges posed by unlabeled and multivariate EEG data. These results establish a new benchmark for future research in hyperscanning studies.

### 4.1. ASC Diagnosis via Deep Learning in Hyperscanning EEG

Despite the growing application of deep learning in EEG studies, its use in diagnosing ASC remains relatively underexplored compared to traditional methods. As noted by Khodatars et al. (2021), neuroimaging modalities such as EEG, MRI, resting-state fMRI, and fNIRS have been widely employed in ASC research, with resting-state fMRI being the most commonly used.

While a handful of studies (Bouallegue et al. (2020); Tawhid et al. (2021); Ranjani and Supraja (2021); Dong et al. (2021) have applied deep learning techniques to EEG data for ASC detection, they primarily focus on individual brain activity. Despite demonstrating promising classification accuracy, these approaches overlook the dynamics of social interactions, which are crucial for understanding ASC. In contrast to these studies, our work shifts the focus to interactions between individuals in hyperscanning, where we analyze simultaneous brain activity from two individuals. This novel approach provides a fresh perspective on ASC detection, offering new insights into how interaction-based neural dynamics can contribute to understanding and diagnosing neurodevelopmental conditions like autism. By reframing psychiatric conditions as emergent from dynamic interpersonal interactions rather than solely individual deficits, we underscore the importance of studying relational processes and mismatched expectations in understanding autism and other conditions (Bolis et al., 2022).

### 4.2. Model Performance and Implications for Clinical Practice

The SSL multi-brain model achieved a mean test accuracy of 78.13%, closely matching the benchmark of 79.4% reported by Banville et al. (2021) for self-supervised learning in EEG classification. This is particularly relevant for advancing diagnostic tools for ASC based on EEG data. Compared to traditional diagnostic tools like the Autism Spectrum Quotient (AQ), Ritvo Autism Asperger’s Diagnostic Scale-Revised (RAADS-R), and the Autism Diagnostic Observation Schedule (ADOS), the model shows both strengths and limitations, in some cases improving upon existing methods.

The model’s sensitivity of 98.90% significantly surpasses the AQ (77%), RAADS-R (52–65%), and ADOS (∼65%) (Ashwood et al., 2016; Conner et al., 2019; Jones et al., 2021; Sizoo et al., 2015). This high sensitivity is key for minimizing false negatives and detecting nearly all ASC interactions, which is crucial for early intervention and improved outcomes. The model’s ROC AUC of 0.8926 also indicates strong discriminative power, outperforming the AQ (0.40) and RAADS-R (0.58), and aligning more closely with the ADOS (0.69). This demonstrates the model’s ability to differentiate ASC behaviors from neurotypical ones.

However, the model faces challenges in specificity, with a rate of 56.47%, leading to a higher rate of false positives. This may be due to overlapping behaviors between neurotypical individuals and those with ASC, a challenge also seen in traditional diagnostic tools like AQ and RAADS-R, where comorbidities such as generalized anxiety disorder (GAD) contribute to false positives (Ashwood et al., 2016; Conner et al., 2019; Jones et al., 2021). The AQ, for instance, has poor specificity (29–52%) in clinical settings due to shared traits with GAD (Ashwood et al., 2016), while RAADS-R’s specificity ranges from 44% to 73% (Conner et al., 2019; Sizoo et al., 2015).

Although not yet ready for widespread clinical adoption, the model has potential as a complementary tool for ASC diagnosis. Its scalability, low-cost post-training, and minimal memory requirements make it suitable for integration into routine clinical practice. The model’s high sensitivity and strong AUC make it particularly effective in identifying ASC interactions. However, further research is needed to improve its specificity, particularly in distinguishing ASC from other conditions with overlapping traits. Ultimately, this study highlights the potential of computational models like the multi-brain approach to enhance diagnostic accuracy, leading to more personalized care for individuals on the autism spectrum.

### 4.3. Modeling Hyperparameters

#### 4.3.1. Epoch Length

In our experimental design, EEG recordings were segmented into one-second epochs as a foundational preprocessing step. This epoch length defined the window size for subsequent pretext tasks, balancing temporal resolution with computational feasibility. Selecting the optimal epoch length is informed by domain knowledge and further refined through experimental tuning. Epoch lengths of one to two seconds are commonly used, as they strike a balance between temporal resolution and statistical reliability (Fraschini et al., 2016; Li et al., 2018). However, the ideal length depends on the study’s objectives and the EEG signal characteristics.

Expanding the size of epochs introduces several trade-offs. While larger epochs may encompass more temporal context, they also reduce the total number of available training samples, increase susceptibility to noise, and exacerbate computational demands due to heightened input dimensionality. Additionally, the artifact rejection pipeline necessitated excluding entire epochs if artifacts were detected, disproportionately affecting longer epochs due to their greater likelihood of containing noise.

Figure 6A illustrates this trade-off using a sample from the HBN dataset. As shown, 1-second epochs led to 28 dropped epochs due to artifacts, while 2-second and 3-second epochs resulted in 35 and 64 dropped epochs, respectively. In contrast, 10-second epochs dropped 60 epochs and caused a marked degradation in signal quality, producing noisier samples. A similar pattern was observed in other samples from both the BBC2 and HBN datasets. However, to conclusively determine the optimal balance between epoch length, artifact rejection rates, and signal quality, further studies are required.

**Figure 6.**
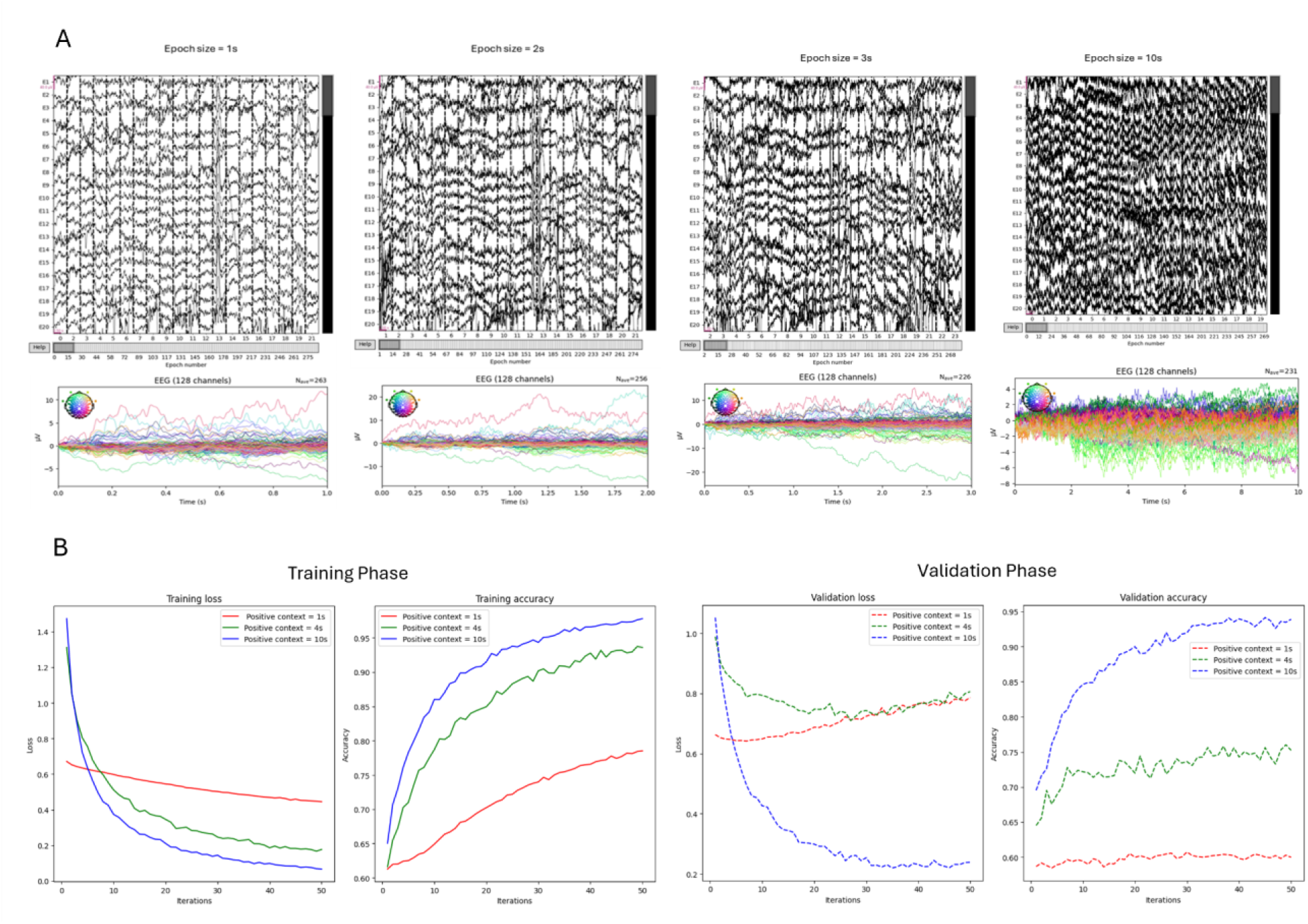
A) Epoch and butterfly plots for each epoch length (sampled from HBN). B) Impact of positive context length in performance of Shallow ConvNet during pretext task training and validation phases.

#### 4.3.2. Positive Context Length

Positive context length, within the pretext task framework, refers to the temporal separation between randomly selected positive epochs and anchor epochs. It is a crucial parameter in our SSL pipeline, influencing the balance of positive and negative samples used for training DL models. A larger positive context length increases the number of epochs labeled as positive, which improves dataset balance. However, if the temporal distance between positive samples becomes too large, the model may struggle to distinguish them effectively, leading to misclassifications. Therefore, selecting an optimal positive context length is essential to balancing dataset distribution and model performance.

To assess its impact, we experimented with positive context lengths of one second, four seconds, and 10 seconds using Shallow ConvNet in the pretext task. As shown in Figure 6B, training loss consistently decreased, and accuracy increased across all context lengths. However, with a one-second positive context, the improvements were slower compared to the four-second and 10-second contexts. During the validation phase, the effect was more pronounced: for the one-second context, loss increased, and accuracy stagnated, indicating difficulties in learning from the minority class and a bias toward the majority class.

Increasing the positive context to four seconds alleviated some of these issues, but performance still showed room for improvement. Notably, extending the context to 10 seconds significantly improved performance, resolving class imbalance and enhancing accuracy. Based on these findings, we selected a positive context length of 10 seconds.

Given our focus on improving downstream task performance, we have not yet explored higher values for positive context length or conducted further hyperparameter tuning. However, as shown, positive context length plays a significant role in the pretext task model’s performance, and further analyses of this parameter are essential. Specifically, investigating the upper bound for positive context length and comparing it to the total duration of EEG recordings could offer valuable insights into optimizing the pretext task model for this dataset and task.

### 4.4. Limitations

This study is subject to several key limitations, which should be considered when interpreting the findings and planning future research.

First, due to the computational demands of training neural networks on large EEG datasets, we fixed the hyperparameters (e.g., learning rate, batch size, number of iterations, dropout rate) for the pretext task. While this decision was made for practical reasons within the scope of the study, it may limit the models’ adaptability to different datasets and training conditions. Future research should explore hyperparameter tuning for the pretext task and assess its impact on downstream model performance. Such exploration could optimize the models for specific tasks and datasets, potentially leading to improved results.

Second, the sampling of epochs in constructing our dataset, as discussed in Sections 4.3.1 and 4.3.2, is influenced by several hyperparameters that significantly affect model performance. While we have observed the effects of positive context length and epoch size, a comprehensive investigation of various hyperparameter combinations is warranted. This would not only enhance model performance but also provide insights into how epoch size, context length, class imbalance, and noise interact to affect model efficacy.

Third, in the logistic regression baseline analysis, we only used 10 of biomarkers proposed by Engemann and colleagues (2018) (section 2.6.2), which were originally designed for single-brain EEG. Our approach to adapting these biomarkers for hyperscanning EEG was simplified: we concatenated individual EEG feature vectors to create a dyadic input. However, this method may not fully capture the complexity of hyperscanning data. Future studies should explore more advanced feature engineering techniques to better adapt single-brain EEG biomarkers to the multi-brain context.

Finally, the models in this study were trained on a subset of the HBN dataset, consisting of 1,000 participants—approximately one-third of the available data. While the HBN dataset is continuously expanding and future versions will include more participants, training on a smaller portion of the data may limit the model’s generalizability. Future studies should address the challenges associated with training on larger datasets, such as optimizing the data loading pipeline to better handle memory constraints for efficient deployment. However, it is also important to investigate the relationship between model complexity, architecture, and dataset size. Understanding this relationship will provide insights into how large the dataset needs to be to achieve optimal performance, and whether further increases in dataset size will continue to yield improvements.

## 5. Conclusion

This study demonstrated the effectiveness of self-supervised learning for classifying hyperscanning-EEG data, specifically in distinguishing between autistic and neurotypical individuals. The SSL multi-brain model significantly outperformed supervised and traditional machine learning baselines, underscoring its ability to extract meaningful patterns from limited labeled data. Our findings highlight SSL’s potential to enhance diagnostic tools for autism spectrum conditions, providing objective and precise assessments that complement traditional methods. This research also emphasizes the broader role of computational models in precision psychiatry, paving the way for innovative, personalized diagnostic and therapeutic solutions.

## Appendix A. HBN and BBC2 Alignment Table

| BBC2 | HBN | BBC2 | HBN | BBC2 | HBN | BBC2 | HBN |
| --- | --- | --- | --- | --- | --- | --- | --- |
| TP10 | E44 | P7 | E58 | P5 | E86 | P3 | E52 |
| P1 | E60 | Pz | E62 | P4 | E92 | P6 | E97 |
| P8 | E96 | PO9 | E68 | PO7 | E65 | PO3 | E67 |
| POz | E72 | PO4 | E77 | PO8 | E90 | PO10 | E94 |
| O1 | E70 | Oz | E75 | O2 | E83 | AF7 | E26 |
| AF3 | E23 | AF4 | E3 | F7 | E33 | F5 | E27 |
| F3 | E24 | F1 | E19 | Fz | E11 | F2 | E4 |
| F4 | E124 | F6 | E123 | F8 | E122 | FT9 | E38 |
| FT7 | E34 | FC5 | E28 | FC3 | E29 | FC1 | E13 |
| FC2 | E112 | FC4 | E111 | FC6 | E117 | FT8 | E116 |
| FT10 | E121 | T7 | E45 | C5 | E41 | C3 | E36 |
| C1 | E30 | Cz | E129 | C2 | E105 | C4 | E104 |
| C6 | E103 | TP8 | E102 | T8 | E108 | TP9 | E57 |
| TP7 | E46 | CP5 | E47 | CP3 | E42 | CP1 | E37 |
| CPz | E55 | CP2 | E87 | CP4 | E93 | CP6 | E98 |

